# Resolving phylogenetic relationships within the *Trichophyton mentagrophytes* complex: a RADseq genomic approach challenges status of “terbinafine-resistant” *Trichophyton indotineae* as distinct species

**DOI:** 10.1101/2024.12.03.626654

**Authors:** Michaela Švarcová, Miroslav Kolařík, Yuanjie Li, Clement Kin Ming Tsui, Vít Hubka

**Affiliations:** Department of Genetics and Microbiology, Faculty of Science, Charles University, Prague, Czech Republic; Laboratory of Fungal Genetics and Metabolism, Institute of Microbiology, Czech Academy of Sciences, Prague, Czech Republic; State Key Laboratory of Mycology, Institute of Microbiology, Chinese Academy of Sciences, Beijing, China; Infectious Diseases Research Laboratory, National Centre for Infectious Diseases, Tan Tock Seng Hospital, Singapore; LKC School of Medicine, Nanyang Technological University, Singapore; Faculty of Medicine, University of British Columbia, Vancouver, Canada; Department of Botany, Faculty of Science, Charles University, Prague, Czech Republic

**Keywords:** anthropophilic dermatophytes, antifungal resistance, dermatomycosis, population structure, taxonomy, *Trichophyton mentagrophytes*, *Trichophyton interdigitale*, zoophilic dermatophytes

## Abstract

The *Trichophyton* (*T.*) *mentagrophytes* complex encompasses common dermatophytes causing superficial mycoses in humans and animals. The taxonomy of the complex is unstable, with conflicting views on the species status of some taxa, particularly *T. indotineae* and *T. interdigitale*. Due to presence of intermediate genotypes, MALDI-TOF MS and ITS rRNA sequencing cannot accurately distinguish all taxa in the complex, potentially contributing to clinical misdiagnoses. In order to resolve the phylogenetic relationship of *T. mentagrophytes* complex, we employed Restriction Site Associated DNA Sequencing (RADseq) to produce a high-resolution single nucleotide polymorphism (SNP) dataset from 95 isolates. The SNP-based analyses indicated the presence of two major genetic clusters, corresponding to *T. mentagrophytes* (including *T. indotineae*) and *T. interdigitale*. Our results challenge the species status of *T. indotineae*, because of insufficient genetic divergence from *T. mentagrophytes*. Therefore, we propose designating *T. indotineae* as *T. mentagrophytes* var. *indotineae* (or *T. mentagrophytes* ITS genotype VIII) to avoid further splitting of the complex and taxonomic inflation. Although *T. interdigitale* shows clearer genetic differentiation, its separation is incomplete and identification of some isolates is ambiguous when using routine methods, leading us to consider it a variety as well: *T. mentagrophytes* var. *interdigitale*. We recommend using T. mentagrophytes as the overarching species name for all complex isolates. Where precise molecular identification is possible, the use of variety ranks is encouraged. Since identical resistance mechanisms are not specific to any genotype or dermatophytes species, identifying antifungal resistance is more important than differentiating closely related genotypes or populations.

**Importance:** Accurate identification of the causal agent is essential for managing dermatophytosis. However, the taxonomy of the *T. mentagrophytes* complex has been plagued by inconsistencies and misidentifications. Our high-resolution SNP analysis revealed that *T. indotineae*, often associated with terbinafine resistance, is not genetically distinct enough from *T. mentagrophytes* to warrant species status. This finding has significant implications for both taxonomy and clinical practice. We suggest consolidating the taxonomy under *T. mentagrophytes*, with variety ranks for *T. indotineae* and *T. interdigitale*. This simplified system will reduce confusion and ensure the comparability of historical and emerging epidemiological data under the umbrella name of *T. mentagrophytes*. By shifting the focus to antifungal resistance detection rather than complex taxonomic distinctions, this reclassification will streamline clinical decision-making. Notably, identical resistance mechanisms are not exclusive to ITS genotype VIII (*T. indotineae*) but also occur in other genotypes and dermatophyte species, albeit with varying prevalence.

## Introduction

*Trichophyton mentagrophytes* is a key pathogen within the diverse dermatophyte group, responsible for causing superficial mycoses affecting skin, nails, and hair in humans and animals. These infections pose a clinical concern of the global population with a high prevalence (1–3). The complexity and variety within *T. mentagrophytes* complex underscore the challenges in dermatophyte taxonomy, necessitating precise species identification for effective clinical management and introduction of preventive measures against the spread of dermatophytoses.

Routine identification of dermatophytes in clinical setting still often relies on morphological characters, or molecular methods such as sequencing of ITS rDNA region or MALDI-TOF mass spectrometry. But in some cases, these methods may falter when discerning closely related dermatophyte species. Such limitations can lead to misdiagnosis and suboptimal treatment, highlighting the need for a more robust understanding of dermatophyte genetics and taxonomy (3–5).

Historically, the species delimitation within the *T. mentagrophytes* complex has undergone many revisions with a rich tapestry of renaming and reclassification efforts (5–9). Yet the most recent taxonomic study (5) recommends to treat *T. mentagrophytes* as a single species with three varieties, including *T. mentagrophytes* var. *mentagrophytes*, *T. mentagrophytes* var. *interdigitale*, and *T. mentagrophytes* var. *indotineae*. Based on this study, there are few taxonomic arguments for preserving these populations as separate species names due to the lack of monophyly and unique morphology among others. This study also challenges the distinctiveness of *T. indotineae*, a widely used name in clinical practice in recent years, known for its association with an Indian dermatophytosis epidemic and a propensity for terbinafine resistance (10–12). Thus, *T. mentagrophytes* complex, presents a substantial challenge in species identification and clinical management due to its diverse morphology, genetic variability, clinical presentation and spread of resistance.

Our study addresses the pressing question of taxonomic classification and population genetic relatedness within the *T. mentagrophytes* complex. As traditional methods such as ITS barcoding, MLST approaches, morphological analysis, and biochemical testing have limitations to conclusively resolve these complex questions, we used Restriction site Associated DNA sequencing (RADseq), a powerful tool that provides comprehensive SNP data and insights into genetic variability and evolutionary relationships. This cost-effective and powerful approach of producing comprehensive SNP (single nucleotide polymorphism) data across the entire genome has been extensively employed in diverse organisms (13–18) but has never been used in dermatophytes. Our findings based on the high-resolution SNP dataset offer a comprehensive re-evaluation of species boundaries, particularly challenging the distinct species status of *T. indotineae*, and propose a simplified and cohesive taxonomic framework.

## Material and Methods

### Isolates

The examined strains originated from Czech patients with various manifestation of dermatophytosis or from foreign culture collections. These strains were previously identified using the DNA sequencing of the ITS rDNA region, translation elongation factor 1-α (*tef1-α*) and β-tubulin gene (*tubb*) in the study of Švarcová *et al.* (2023). A subset of 95 isolates examined in this study represents all major clades identified in *T. mentagrophytes* complex from the aforementioned research.

### DNA Extraction

Cultures were grown in liquid Sabouraud’s glucose medium (Thermo Fisher Scientific, Waltham, MA, USA), shaking at 100 rpm for 10 days. DNA was extracted using the QuickDNA Miniprep Kit (Zymo Research, Irvin, CA), following the manufacturer’s protocol. The quality and quantity of the DNA were measured using Qubit dsDNA HS Assay Kit (Thermo Fisher Scientific, Waltham, MA, USA). The DNA was normalized to a concentration of 20 ng/µL.

### Sequencing, Variant calling and genome assembly

Library preparation, enzyme selection and RAD sequencing reaction were performed by Floragenex Inc (9590 SW Gemini Dr, Beaverton, OR, USA). The PstI enzyme was used for digestion. From a total of 95 samples, 409.8 million reads were obtained, achieving an average coverage of 20,187.5 × per variant. This allowed to identify 16,795 variable loci. The reference genome for strain ME 517/15 was assembled using VELVET software v. 1.2.10 (19). The remaining samples were aligned to this reference genome and processed into variant call format (vcf) using the BOWTIE v. 1.1.1 (20), BWA v. 0.6.1 (21), and SAMTOOLS v. 0.1.16 (22).

### Population structure

The vcf file generated above was used for SNP data processing. TASSEL v. 5.2.91 (23) was used for quality control, with initial filtering removing taxa with more than 10% missing sites, reducing the dataset from 95 into 87 individuals. Further filtering eliminated missing sites, parameter Site Minimum Count of alleles not Unknown was set to 80, and Site Minimum Minor Allele Frequency to 0.1 (10%), reducing the number of sites from 16,795 to 6,996.

For performing population structure analysis and SNP phylogeny analysis, only individuals with unique haplotype were included (n = 18). The population structure analysis was performed using the set of scripts STRUCTURE multi-PBS Pro (24), which enables parallel execution of the STRUCTURE software on a high-performance computing (HPC) cluster using the Portable Batch System (PBS) Pro job scheduler. This method allows for efficient and simultaneous analysis of multiple datasets or multiple runs of the STRUCTURE program (25). The analysis spanned a range of clustering number (K) values from 1 to 10, with each subset run ten times using 1,000,000 replicates and burn-in 500,000. Computational resources were provided by MetaCentrum. The Evanno ΔK statistic was calculated using STRUCTURE HARVESTER (26). Filtered SNPs were also used to construct a TCS haplotype network using the PopART program (27).

### Discriminant Analysis of Principal Components (DAPC)

Also, the genetic structure among the isolates was investigated using DAPC. First, the vcf file containing the variant information was imported using vcfR and converted into a genlight object (28). Second, the find.cluster function in the R package adegenet v 2.0.1 was used to identify clusters, and the optimal K was determined by the lowest value of Bayesian Information Criterion (BIC) (29). We set max.n.clust = 40 and n.iter = 1 × 10^9^. The optimal number of principal components was determined by ‘a score’ using the optim.a.score function in the adegenet package. The final DAPC was performed using the optimal K with the best score, and the scatter plot was plotted using ggplot2 v3.3.5 (30).

### Phylogeny

SNAPP v. 1.6.1 (31), an add-on package to BEAST 2 software (32), was utilized for SNP tree computation. The Markov Chain Monte Carlo (MCMC) chain length was set to 10^7^, and samples were collected every 1000 iterations. The results obtained from the SNAPP analysis indicated that the effective sample size (ESS) for the parameters estimated by the MCMC chains was greater than 200, suggesting effective convergence. The log files were reviewed using Tracer to ensure the accuracy and reliability of the analysis. To visualize the posterior distribution of species trees, which represents the most likely tree topology, we utilized the DensiTree v. 2.2.7. This software overlays all the sampled tree topologies onto a single plot, providing a visual representation of the uncertainty and variation in the estimated species tree. A total of 10,000 trees were processed using TreeAnnotator v. 2.7.4, with the initial 20% of trees discarded. The consensus topology was generated from the remaining 8,000 trees and visualized using Figtree v1.4.4 (33).

An ITS rDNA-based tree was constructed for comparison with phylogeny based on SNP data. To determine the partitioning schemes and suitable substitution models (Bayesian Information Criterion) , PartitionFinder 2 was used (34). The ITS region was segmented to ITS1, 5.8S and ITS2 regions which were considered as independent datasets. The optimal partitioning schemes identified for the dataset were TrNef+I for ITS1 and ITS2 regions, and JC 5.8S region. A maximum likelihood (ML) phylogenetic tree was constructed in IQ-TREE version 2.1.2 (35), with nodal support assessed through nonparametric bootstrapping (BS) with 1000 replicates. The comparison of two phylogenetic trees was visualized using a tanglegram, with the samples connected by lines based on their names. The tanglegram visualization was generated using R Studio (36) and R packages ape and phytools (37, 38).

## Data availability

The filtered dbSNP VCF dataset used is available through DRYAD digital repository https://doi.org/10.5061/dryad.zpc866tjn (Link for reviewers: https://datadryad.org/stash/share/rotYt9h27LckCoxj7YxYlnN8JC_mmDu06UMvNWU <u>t_qQ</u>). Selected isolates used in this study are available in the CCF collection of fungi (Charles University, Prague, Czechia).

## Results

### Population structure

Analysis of the population structure using Evanno ΔK statistics revealed that the scenario ΔK = 2 was the most probable, while the scenario ΔK = 4 was the second most likely. This suggests that our dataset can be classified into either two or, less likely, four distinct populations. Fig. 1 illustrates these findings, showing the Evanno ΔK statistics graph (Fig. 1B) alongside pophelper-generated bar plots from STRUCTURE runs for ΔK = 2, ΔK = 3, and ΔK = 4 (Fig. 1A).

**FIG 1.**
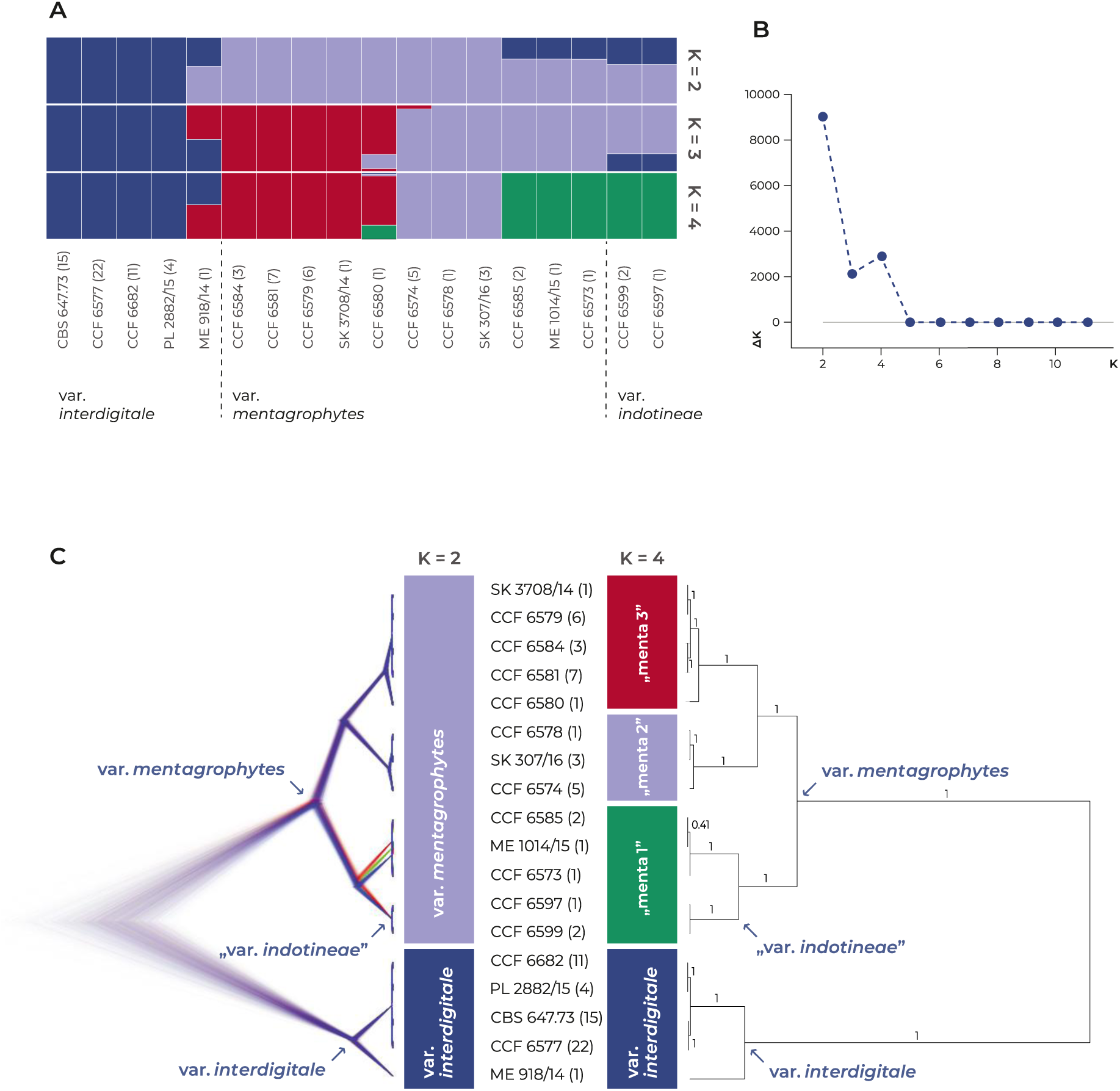
Population structure and phylogenetic relationships of populations within the *Trichophyton mentagrophytes* complex based on SNP data. (**A**) Bar plots displaying the population structure based on STRUCTURE analysis, showing clustering for K=2, K=3, and K=4. Each vertical bar represents a unique haplotype (the numbers in brackets indicates the number of identical strains), with colors corresponding to the proportion of ancestry from each inferred population. (**B**) The Evanno ΔK statistic, indicating that the K=2 is the most likely scenario, while K=4 represents the second most probable scenario. (**C**) Phylogenetic tree generated using BEAST based on SNP data. The left side displays the BEAST tree visualized using DensiTree, trees with the most common topology are highlighted by blue, trees with the second most common topology by red, trees with the third most common topology by green. On the right side, the consensus phylogenetic tree (posterior probabilities supporting branches are appended to nodes) is shown. Populations identified in the STRUCTURE in scenarios K=2 and K=4 are indicated by colored bars in the middle.

In the most probable scenario (ΔK = 2), the isolates were distributed into two population corresponding to *T. mentagrophytes* var. *mentagrophytes* and *T. mentagrophytes* var. *interdigitale*, as previously identified using multi-locus sequence typing (MLST) (5). *Trichophyton mentagrophytes* var. *indotineae* was not recognized in this scenario and was included in *T. mentagrophytes* var. *mentagrophytes*. However, significant number of haplotypes showed high level of admixture between these two populations indicating possible recombination and incomplete separation of *T. mentagrophytes* var. *mentagrophytes* and *T. mentagrophytes* var. *interdigitale*.

The second most probable scenario (ΔK = 4) recognizes *T. mentagrophytes* var. *interdigitale* as a distinct group and divides *T. mentagrophytes* var. *mentagrophytes* into three groups: menta1 (green), menta2 (pale blue), and menta3 (red) (Fig. 1). *Trichophyton mentagrophytes* var. *indotineae* was again not recognized as a separate population and was included in group menta1 of *T. mentagrophytes* var. *mentagrophytes* along with some other isolates not belonging to *T. mentagrophytes* var. *indotineae* (ITS genotype VIII). There was a significant level of admixture between *T. mentagrophytes* var. *interdigitale* and population menta3, as seen in example of strain ME 918/14, and also between populations menta1 and menta3 as seen in example of strain CCF 6580. This indicates that these populations are not fully separated.

### Phylogeny and haplotype network

A total of 87 isolates meeting the quality control (QC) criteria were selected for analysis. We initially constructed a Bayesian tree using BEAST 2 based on 6,996 SNPs extracted from examined strains that remained after QC and filtering (see Methods). Fig. 1C presents a DensiTree visualization of consensus species tree, showcasing the confidence level across various trees topologies. The Consensus topology tree, depicted to the right in Fig. 1C, highlights the consensus topology across sampled trees. Predominantly high posterior probabilities (pp) denote strong support (mostly pp = 1.00), with only one branch having pp of 0.41, reflecting poorly resolved relationships between some members of the menta1 population. The colored bars inserted between the trees show corresponding distribution of populations according to the STRUCTURE analysis (most probable scenarios ΔK = 2, and ΔK = 4).

The haplotype network (PopART) and unrooted SNP-based tree (BEAST 2) illustrated in Fig. 2 provide detailed insights into the relationships among *T. mentagrophytes* complex strains and populations. The unrooted phylogenetic tree illustrates the phylogenetic distances between isolates and clearly distinguishes *T. mentagrophytes* var. *interdigitale* from *T. mentagrophytes* var. *mentagrophytes*, highlighting their relatively high genetic divergence compared to divergencies between subpopulations present within *T. mentagrophytes* var. *mentagrophytes*. However, *T. mentagrophytes* var. *indotineae* is included within *T. mentagrophytes* var. *mentagrophytes* (scenario ΔK = 2) or within its subpopulation menta1 (scenario ΔK = 4).

**FIG 2.**
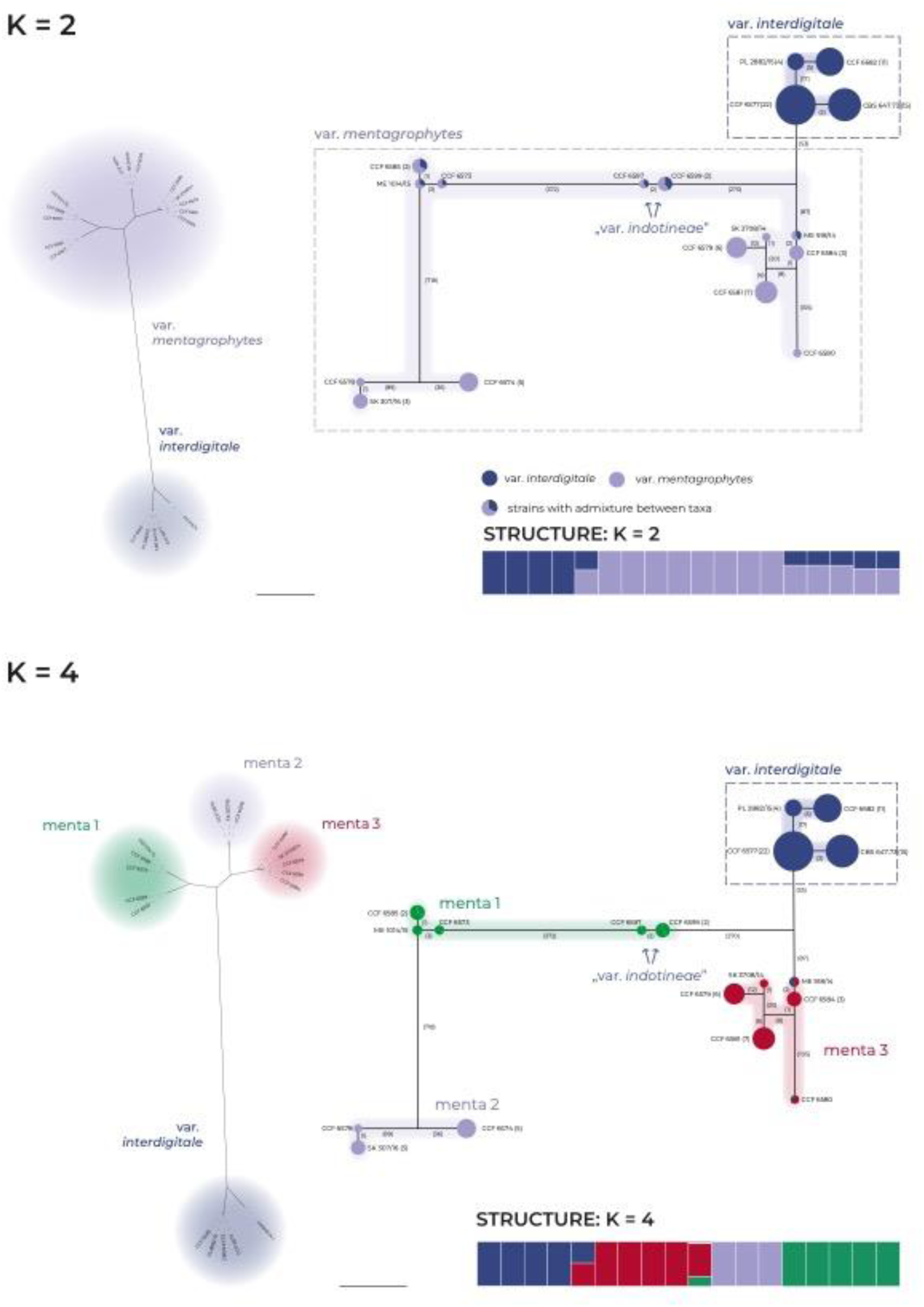
Clustering patterns of the *Trichophyton mentagrophytes* complex based on SNP data. The figure shows population structure and genetic relationships for K=2 and K=4 scenarios inferred from STRUCTURE. The relationships are illustrated on haplotype network (right parts of panel, PopART software) and unrooted phylogenetic tree (left parts of panel, BEAST 2 software). In the haplotype network, the haplotypes are represented by circles with size corresponding to number of isolates with identical haplotype, numbers on connecting lines indicate number of substitutions between haplotypes (indels excluded), the colors represent the clusters from STRUCTURE analysis, presence of a multiple colors in one circle indicates admixture for a given haplotype and scenario (bar plot below network; for details see Fig. 1). The unrooted phylogenetic trees based on SNPs (left parts of panels), depicting the phylogenetic distances among the unique haplotypes and their affiliation to clusters from STRUCTURE analysis.

### Comparison of SNP-based and ITS-based phylogenies

There is a good congruence between ITS genotypes and SNP tree (Fig. 3). In the SNP-based phylogeny, the populations of *T. mentagrophytes* var. *interdigitale* and *T. mentagrophytes* var. *mentagrophytes* are well separated and monophyletic, while in the ITS tree, they are paraphyletic. In both phylogenies, isolates of *T. mentagrophytes* var. *indotineae* are placed within *T. mentagrophytes* var. *mentagrophytes* (subpopulation menta1). In the SNP-based phylogeny, the isolate CCF 6580 is part of subpopulation menta3 of *T. mentagrophytes*, whereas the ITS phylogeny positions this isolate separately from other *T. mentagrophytes* strains.

**FIG 3.**
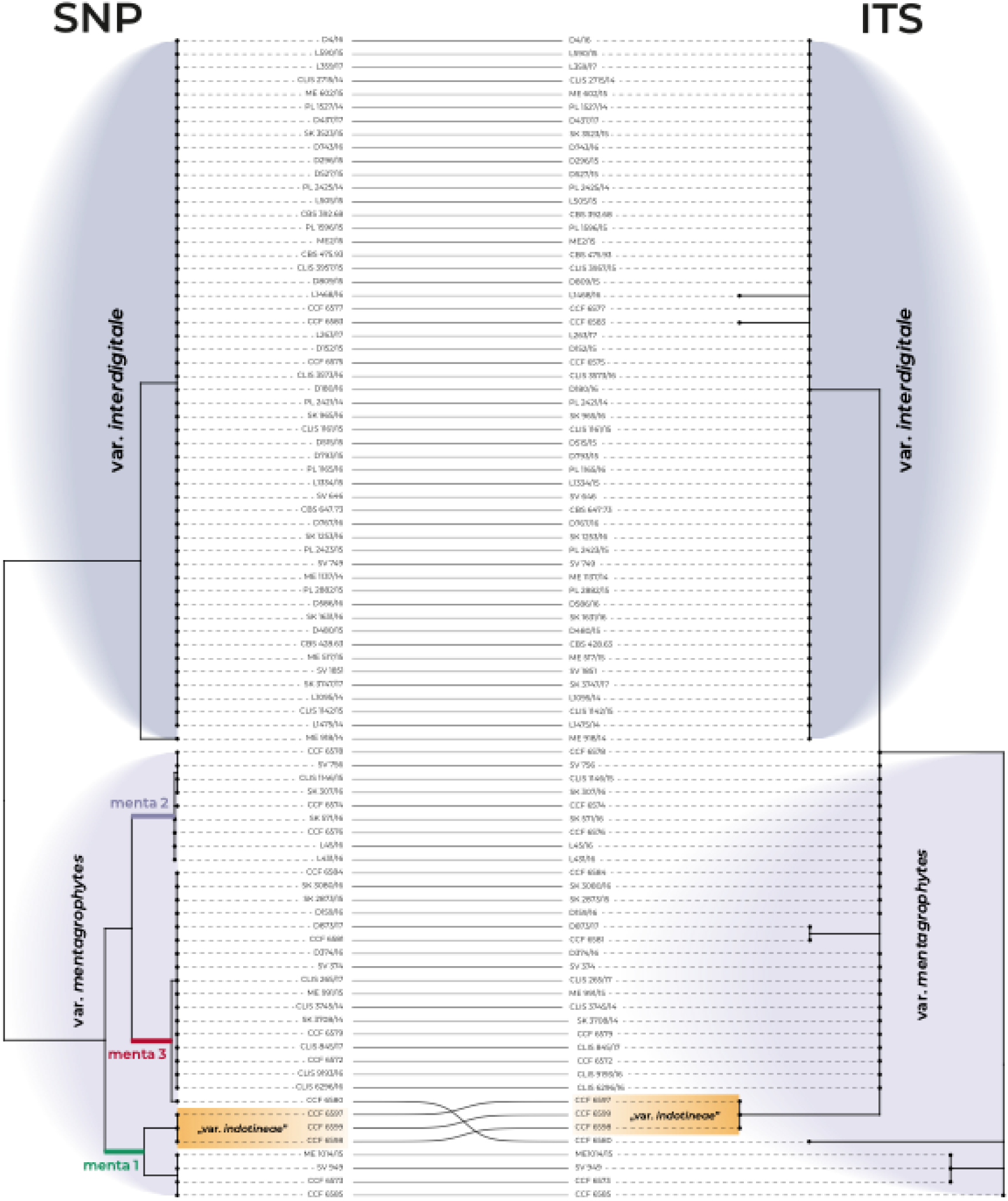
Comparison of phylogenetic trees topologies based on SNP data (RADseq) and ITS region within the *Trichophyton mentagrophytes* complex, connected by tanglegram. Lines connect corresponding isolates between the trees for visual clarity. The SNP-based phylogeny provides better resolution within *T. mentagrophytes* var. *mentagrophytes*, identifying three main subpopulations (menta1, menta2 and menta3). Both trees show low resolution within *T. mentagrophytes* var. *interdigitale*, which is resolved as monophyletic in the SNP-based tree but paraphyletic in the ITS-based tree. Additionally, *T. mentagrophytes* var. *indotineae* is integrated within *T. mentagrophytes* var. *mentagrophytes* in both analyses.

### DAPC

To gain insights into the population structure of *T. mentagrophytes* complex, we performed DAPC analysis using SNPs markers. DAPC plots revealed high level of population differentiation, and relatively consistently clustered samples that correspond to the *T. mentagrophytes* var. *interdigitale*, as well as menta1, menta2 and menta3 (Fig. 4). *Trichophyton mentagrophytes* var. *indotineae* as currently circumscribed was differentiated from other four strains of menta1 subpopulation. Notably, isolates CCF 6580 (menta3) and ME 918/14 (var. *interdigitale*) were clearly distinct from the remaining strains in the respective populations, potentially representing genotypes or populations that require additional sampling.

**FIG 4.**
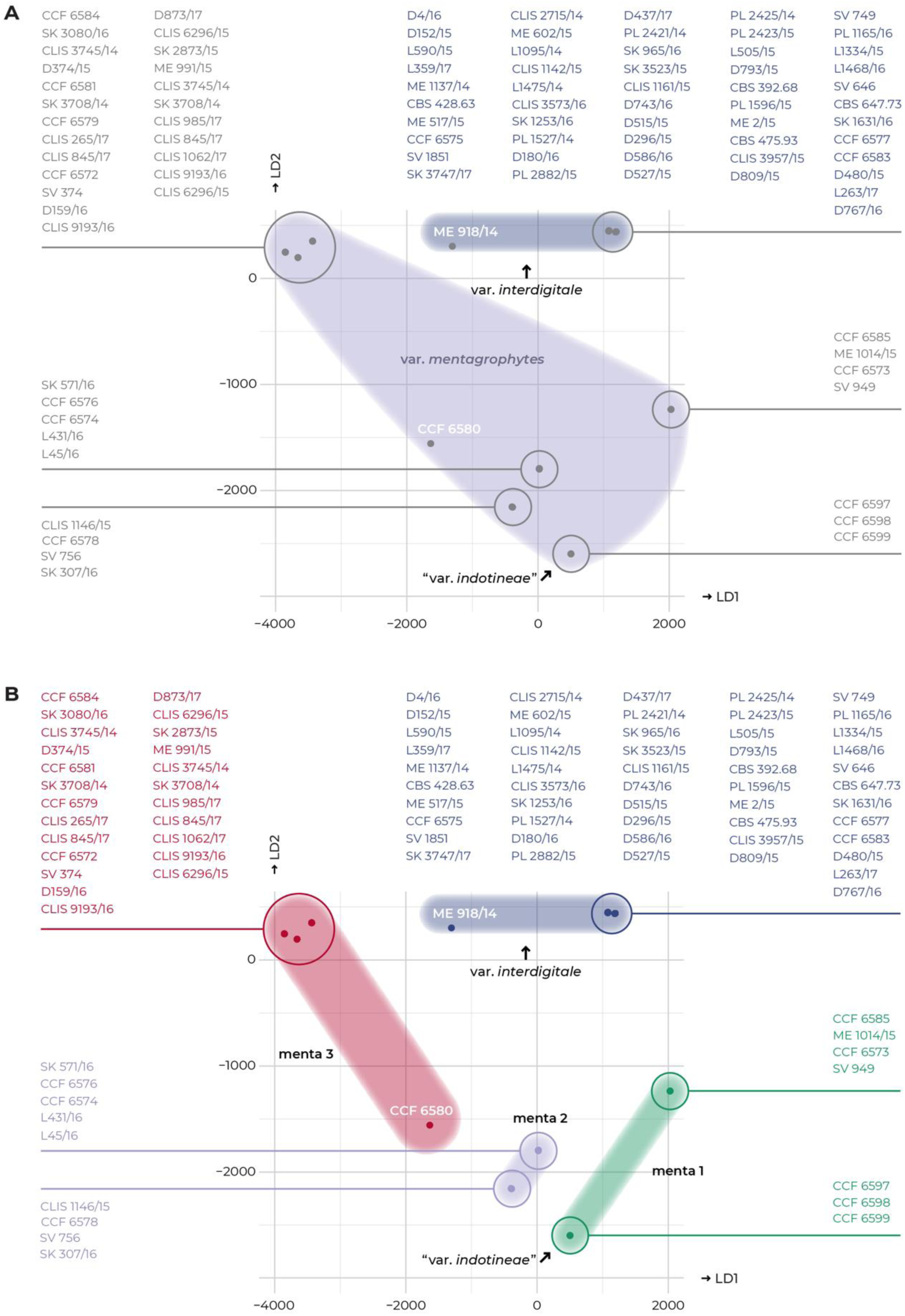
Discriminant analysis of principal components (DAPC) based on SNP data showing clustering of strains within *Trichophyton mentagrophytes* complex. Colored clouds represent clustering from STRUCTURE analysis with K = 2 (A) and K = 4 (B). The analysis shows clustering of strains. The analysis does not show clear clustering of isolates based on taxonomic varieties (var. *mentagrophytes*, var. *interdigitale*, and var. *indotineae*).

## Discussion

This study provides substantial new insights into the taxonomy and species delimitation within the *Trichophyton mentagrophytes* complex using a RADseq genomic approach. These findings challenge the current status of *T. indotineae* as a distinct species, suggesting it should instead be considered a variety or integral part of *T. mentagrophytes*.

The primary goal of this study was to address the taxonomic uncertainties within the *T. mentagrophytes* complex, which has historically been plagued by numerous taxonomic changes and misidentifications (5–8, 39). The methods commonly used for identification, such as phenotype, ITS region sequencing and MALDI-TOF MS, often fail to distinguish related taxa within the complex due to their limitations or the presence of transitional forms and genotypes (39–41). The ambiguous or challenging identification leads to situations where the authors, despite using molecular data for species identification, prefer to use only the designations “*T. mentagrophytes* complex” and “*T. mentagrophytes/T. interdigitale*” to refer to all strains from the complex (42–48).

Our population structure analysis using data from RAD sequencing revealed that the most probable scenario with presence of two populations in the *T. mentagrophytes* complex (ΔK = 2), corresponds to *T. mentagrophytes* (including *T. indotineae*) and *T. interdigitale* (Fig. 1). This finding suggests that *T. indotineae* does not possess sufficient genetic divergence to be considered a separate taxonomic entity. Its recognition at species level could lead to taxonomic inflation and further unnecessary segregation of *T. mentagrophytes* into multiple species to retain monophyly of recognized taxa. In the second most probable scenario with four populations (ΔK = 4), *T. interdigitale* is recognized as a separate population and *T. mentagrophytes* is divided into three populations (menta1, menta2 and menta3), none of which corresponds to *T. indotineae*, reinforcing the notion that it should not be recognized as an independent species.

The use of approaches based on whole genome sequencing (WGS) are still uncommon in dermatophytes and only a few studies performed WGS to address various hypotheses (39, 49–52). From a taxonomic point of view, it is particularly noteworthy the work of Pchelin *et al.* (39) who concluded based on phylogenomic data and species delimitation analyses that *T. mentagrophytes* and *T. interdigitale* belong to the same phylogenetic species. The authors, however, argued that both species should be retained due to the correlation of epidemiological data with ITS genotypes and taxonomic stability. Other well-established method for population analysis in dermatophytes involves in particular microsatellite genotyping and MLST (multilocus sequence typing). The MLST approaches reached the same conclusion that *T. mentagrophytes* and *T. interdigitale* and/or *T. indotineae* are not monophyletic as reviewed by Švarcová et al. (5). Although microsatellite markers were used in many clinically important species complexes, Including *T. rubrum* (53, 54), *T. benhamiae* (55–57) and *Microsporum canis* (58–60), they are not available in *T. mentagrophytes* complex.

### Summary of reasons for not recognizing populations in the complex as separate species

#### 1. Lack of monophyly

Numerous studies performing phylogenies based on ITS region, multiple genes (6, 11, 61) and phylogenomic data (39, 52) demonstrated that taxa in the complex are not monophyletic. Overuse of the rank of a species leads to taxonomic inflation and could lead to subdivision of the *T. mentagrophytes* complex into additional species without biological relevance so that monophyly is maintained. These situations are well known in many fungal general such as *Alternaria* (62), *Aspergillus* (63, 64), *Bryoria* (65), *Diaporthe* (66), *Flammulina* (67) and others, and led to significant species reduction in recent years. The use of taxonomic levels below the species level such as variety, subspecies and others is appropriate for the aforementioned cases when we want to describe biologically or clinically relevant populations (46). For these units, there is no pressure to meet all the usual criteria used to define species under study, including monophyly.

#### 2. Lack of unique morphology

Reliable differentiation of *T. mentagrophytes*, *T. interdigitale* and *T. indotineae* is practically impossible using phenotypic features (5, 10, 11). Although some statistically significant differences were found between these species, there is an important overlap in the presence/absence of features or range of values making these characters unusable for accurate identification in clinical laboratories. For instance, Švarcová et al. (5) demonstrated that *T. interdigitale* strains had significantly slower growth after 7 days at 37 °C (6–24 mm; 15 mm on average) than *T. mentagrophytes* strains (8–32; 22 mm on average). Tang et al. (11) reported statistically significant differences in urease activity between *T. mentagrophytes* (67% strains hydrolysed urea) and *T. interdigitale* (75%) versus *T. indotineae* (27% strains hydrolysed urea). The hair perforation showed similar results with 92.5% of *T. mentagrophytes* and 98.5% of *T. interdigitale* having positive result, while only 27% of *T. indotineae* showed. As is evident, none of these characters are diagnostic for mentioned populations/species.

#### 3. Antifungal resistance and treatment

The treatment of dermatophytosis is guided by the recommendations specific to particular clinical unit (such as tinea capitis, tinea corporis, tinea unguium, etc.), along with an understanding of the local situation regarding resistance levels. Terbinafine resistance, which has been increasingly documented in recent years in the genus *Trichophyton* (68–73), is most commonly due to non-synonymous point mutations in the squalene epoxidase gene (SQLE). The amino acid substitution F397L in SQLE is one of the most commonly observed substitutions worldwide associated with a high level of terbinafine resistance (68). This mechanism of resistance is not specific to any genotype or population in the *T. mentagrophytes* complex. The same mechanism has been described in *T. mentagrophytes* (69), *T. interdigitale* (*74*) and *T. rubrum* (70–72). Although the ITS genotype VIII of *T. mentagrophytes* (*T. indotineae*) is often labeled as “terbinafine-resistant”, its resistance level varies greatly. In other words, resistance is not an intrinsic characteristic of *T. indotineae* and only a portion of its isolates display resistance. Although some studies reported resistance in all strains collected, e.g., study from the Greece (9/9 strains) (75) and North America (21/21 *T. indotineae* strains) (76), the majority of epidemiological studies showed lower levels of resistance. Namely, 16.7 % (1/6) in France (isolates collected in 2021 only) (77) , 45% (13/29) in Germany (2016–2019) (78), 50% (2/4 strains) in Vietnam (2020–2021) (79), 53% (34/64) in India (2018–2019) (80), 72% (202/279) in India (2017–2019) (81), and 78 % (7/9) in the Czech Republic (2018–2022) (69). Identical resistance mechanism is also known in other ITS genotypes of *T. mentagrophytes*, including ITS genotype VII (82), and ITS genotype XVII (73). This demonstrates that arguments advocating the recognition of *T. indotineae* because of its resistance are unfounded, and that antifungal susceptibility testing is superior to species/genotype identification. It is true that the level of resistance in this genotype is higher than in other genotypes and species, likely due to positive selection pressure, which confers a survival advantage to resistant strains, allowing them to spread more effectively because of their resistance to treatment. This scenario can happen in any resistant genotype or population. Species identification as ‘terbinafine-susceptible’ or ‘terbinafine-resistant’ *T. mentagrophytes* provides more clinically meaningful information compared to identification as *’T. indotineae*’ only as strains classified under *T. indotineae* may exhibit both susceptibility and resistance.

Similar situations, where resistance is predominantly associated with a specific subpopulation of a clinically relevant species, can be found in other clinically significant fungi. For example, azole resistance is primarily associated with MLST clade 4 of *Candida tropicalis* (83) and clade A/population A of *Aspergillus fumigatus* (84, 85). Although these subpopulations can be clearly identified using DNA sequencing or subtyping methods, the resistant population has not been reclassified as a new species, unlike *T. indotineae*.

#### 4. Clinical and ecological definition

It has repeatedly been confirmed that onychomycosis and tinea pedis are more commonly linked to *T. interdigitale* than *T. mentagrophytes* (5, 39, 86–89) and patients infected with *T. interdigitale* are significantly older (5) as onychomycosis is more prevalent patients in older age groups. While *T. interdigitale* is usually considered anthropophilic and *T. mentagrophytes* zoophilic dermatophyte, there are some other sublineages/genotypes in *T. mentagrophytes sensu lato* that primarily spread among humans, including ITS genotype VIII (*T. indotineae*) and ITS genotype VII (90). This may partly reflect the lack of research in the veterinary field, as evidenced by isolations of the ITS genotype VIII (91–93) and VII (94, 95) from animals. These findings suggest that *T. mentagrophytes* cannot be easily classified into anthropophilic or zoophilic ecological group, but as a species with a broad host spectrum that includes a wide range of mammals, including humans, with certain lineages more or less specialized to specific hosts. Designating of some clinically relevant lineages as varieties (or genotypes) can raise awareness of these lineages and maintain the integrity of this species as a whole.

Although *T. indotineae* is usually ascribed as an agent of severe cases of tinea corporis, tinea cruris, and tinea faciei (78), some recent studies from Switzerland (96) and Czech Republic (69) revealed that this genotype is most frequently associated with onychomycosis, making it challenging to use the clinical picture as a taxonomic criterion to define *T. indotineae*.

#### 5. Lack of simple identification molecular tools and identification ambiguities

The ability to easily and accurately identify pathogens is crucial in clinical practice. Differentiating *T. mentagrophytes* and *T. interdigitale* using MALDI-TOF, a method that has gained popularity due to its speed and efficiency, usually fails (40, 47, 97). In contrast, MALDI-TOF probably enables the identification of *T. indotineae* from *T. mentagrophytes/T. interdigitale* with an accuracy rate of 96% or higher (40, 47). However, it does not differentiate terbinafine-susceptible and resistant strains (98). A key limitation of these studies is that they analyzed a limited number of strains representing restricted variability within *T. mentagrophytes* (many ITS genotypes were missing). As a result, the identification accuracy of *T. interdigitale* and *T. mentagrophytes* is expected to decline even more in broader studies. The same concern may apply to *T. indotineae* strains, where further limitations could arise when comparing it to other closely related genotypes of *T. mentagrophytes*.

The differentiation of *T. indotineae* from *T. mentagrophytes*/*T. interdigitale* relies only on 1–2 unique substitutions in the ITS region (depending on the alignment length and intraspecific diversity included) (5, 40), and no unique substitutions were found in other commonly used markers such as *tef1-α* and *tubb* (5). This highlights how poorly defined these populations are, aligning with our results based on SNPs obtained through RADseq method, where *T. indotineae* is located inside menta1 subpopulation of *T. mentagrophytes* var. *mentagrophytes*.

The ITS region genotyping, increasingly used to characterize isolates of the *T. mentagrophytes* complex (12, 89, 99, 100), is relatively time-consuming and requires expertise. From practical standpoint, it may be advisable to differentiate ITS genotypes that predominantly spread via human-to-human contact to carry out an epidemiological investigation, i.e., I, II, X, XI, and XII (*T. mentagrophytes* var. *interdigitale*), VII (mostly associated with sexually transimitted dermatophytosis) and VIII (*T. mentagrophytes* var. *indotineae*). However, animal reservoirs cannot be ruled out (discussed above), and ecological classification is further complicated by the existence of intermediate genotypes II* and III* (12, 39, 86, 101) between *T. mentagrophytes* var. *interdigitale* and *T. mentagrophytes* var. *mentagrophytes* that more frequently originate from animals (11, 12, 39, 86).

## Conclusions and recommendations

This study reexamines the taxonomy of the *T. mentagrophytes* complex, particularly challenging the species status of *T. indotineae* and addressing broader taxonomic issues. Our phylogenetic and population-genetic analysis, based on whole-genome SNP data generated through RADseq, revealed insufficient genetic divergence to support the recognition of *T. indotineae* as a distinct species. In contrast to *T. indotineae*, the population of *T. interdigitale* appears better separated from *T. mentagrophytes* based on whole-genome SNP data. Despite this, due to the ambiguous definition of this species and the difficulty of identification by routine methods, we recommend using the name the name *T. mentagrophytes* for all isolates within the complex. When molecular identification of *T. interdigitale* and *T. indotineae* is clear and unambiguous, we suggest optionally assigning them variety ranks: *T. mentagrophytes* var. *interdigitale* and *T. mentagrophytes* var. *indotineae* (alternative: *T. mentagrophytes* ITS genotype VIII). This practice will contribute to cohesive taxonomy, reduce confusion in both clinical and research contexts, and simplify identification efforts in clinical practice. By adopting this taxonomic framework, we ensure that historical and current epidemiological data remain comparable while maintaining clinical distinctions of some subpopulations. This reclassification will also prevent unnecessary taxonomic inflation and streamline diagnostic processes, allowing clinicians to focus on identifying antifungal resistance rather than differentiating genotypes with limited clinical relevance.

## Acknowledgments

The project was supported by the Czech Ministry of Health (grant NU21-05-00681), Strategy AV21 of the Academy of Sciences of the Czech Republic ‘Mycolife program - the world of fungi’, and Czech Academy of Sciences Long-term Research Development Project (RVO: 61388971). We are grateful to Jan Karhan for help with the graphical adjustments of Figures. We thank Soňa Kajzrová and Kateřina Glässnerová for their invaluable assistance in the laboratory. The research reported in this publication was part of the long-term goals of the ISHAM working group Onygenales. Computational resources were provided by the e-INFRA CZ project (ID:90254), supported by the Ministry of Education, Youth and Sports of the Czech Republic.

